# Pharmacological blockade of muscle afferents and perception of effort: a systematic review with meta-analysis

**DOI:** 10.1101/2021.12.23.474027

**Authors:** Maxime Bergevin, James Steele, Marie Payen de la Garanderie, Camille Feral-Basin, Samuele M. Marcora, Pierre Rainville, Jeffrey G. Caron, Benjamin Pageaux

**Affiliations:** École de kinésiologie et des sciences de l’activité physique (EKSAP), Faculté de médecine, Montreal, Canada; Centre de recherche de l’Institut universitaire de gériatrie de Montréal (CRIUGM), Montreal, Canada; School of Sport, Health and Social Sciences, Southampton, United Kingdom; Department of Biomedical and Neuromotor Sciences (DiBiNeM), University of Bologna, Bologna, Italy; Département de stomatologie, Faculté de médecine dentaire, Université de Montréal, Montreal, Canada; Centre de recherche interdisciplinaire en réadaptation du Montréal métropolitain, Montreal, Canada; Centre interdisciplinaire de recherche sur le cerveau et l’apprentissage (CIRCA), Montréal, Canada

## Abstract

**Background:** The perception of effort (PE) provides information on task difficulty and influences physical exercise regulation and human behavior. This perception differs from other-exercise related perceptions such as pain. There is no consensus on the role of group III-IV muscle afferents as a signal processed by the brain to generate PE.

**Objective:** The aim of this meta-analysis was to investigate the effect of pharmacologically blocking muscle afferents on the PE.

**Methods:** Six databases were searched to identify studies measuring the ratings of perceived effort (RPE) during physical exercise, with and without pharmacological blockade of muscle afferents. Articles were coded based on the operational measurement used to distinguish studies in which PE was assessed specifically (*effort dissociated*) or as a composite experience including other exercise-related perceptions (*effort not dissociated*). Articles that did not provide enough information for coding were assigned to the *unclear* group.

**Results:** The *effort dissociated* group (*n*=6) demonstrated a slight RPE increase with reduced muscle afferents feedback (standard mean change raw (SMCR), 0.39; 95%CI, 0.13 to 0.64). The group *effort not dissociated* (*n*=2) did not reveal conclusive results (SMCR, −0.29; 95%CI, −2.39 to 1.8). The group *unclear* (*n*=8) revealed a slight RPE decrease with reduced muscle afferents feedback (SMCR, −0.27; 95%CI, −0.50 to −0.04).

**Conclusions:** The heterogeneity in results between groups reveals that the inclusion of perceptions other than effort in its rating influences the RPE scores reported by the participants. The absence of decreased RPE in the *effort dissociated* group suggests that muscle afferents feedback is not a sensory signal of PE.

**Key points:**

- To date, there is no consensus on the neurophysiological signal processed by the brain to generate the perception of effort.
- Following a systematic search in six databases, this meta-analysis suggests that reducing afferent feedback from the working muscles via epidural anesthesia does not reduce perception of effort.
- This systematic review suggests that afferent feedback from the working muscles is not the neurophysiological signal processed by the brain to generate the perception of effort.

## 1. Introduction

During physical exercise, the perception of effort provides information on how intense and difficult the task being performed is perceived. The perception of effort is involved in the regulation of human behavior [1] and influences how the nervous system selects a given movement amongst a myriad of possibilities [2, 3]. The perception of effort is altered in the presence of fatigue [4] and various pathologies such as chronic fatigue syndrome [5, 6], stroke [7] and cancer [8]. This perception is used to prescribe and monitor exercise in both rehabilitation programs [9, 10] and athletic training [11–13]. Despite the growing interest in this perception, to date, researchers have failed to reach a consensus on the signal(s) processed by the brain leading to its generation [14–16].

One popular model amongst exercise physiologists, referred to as the afferent feedback model, suggests that the feedback originating from the peripheral organs active during physical exercise (i.e., skeletal muscles, heart, lungs) is processed by the central nervous system to generate the perception of effort [17, 18]. Notably, authors suggested that feedback from group III-IV muscle afferents plays an important role the perception of effort [19–21]. The ratings of perceived effort intensity would then be predicted to increase with higher discharge rates of the muscle afferents accompanying intense exercise [22, 23]. In contrast, a popular model amongst neuroscientists and physiologists interested in the regulation of cardiovascular responses during exercise and/or kinesthesia is the corollary discharge model. This model proposes that the perception of effort is generated by the processing of a copy of the central motor command, named the corollary discharge [24–26]. In this model, an increase in the magnitude of the central motor command should result in an increase in the perception of effort intensity [24, 27]. It is important to note that this model does not bar peripheral contributions to the regulation of central commands during voluntary movement, but states that central processing of afferent feedback does not generate the perception of effort and that effort could be perceived in the absence of afferent feedback [14, 15, 28, 29]. For example, any mechanisms able to alter the muscle force production capacity [24, 30] or the corticospinal excitability – changes in cortical and/or spinal excitability influencing the ease with which the central motor command is relayed to working muscles [31] – may modulate the perception of effort by increasing or decreasing the magnitude of the central motor command needed to sustain a given level of performance [32]. For instance, neuromuscular fatigue [24, 27] and pain [33, 34] have both been suggested to affect muscle force production capacity and corticospinal excitability. As such, in the presence of both phenomena, an increase of the central motor command might be required to recruit additional motor units and keep the same level of performance, thereby increasing perceived effort.

A powerful technique to gain insights into the neurophysiology of the perception of effort and test the existing models is by pharmacologically blocking group III-IV muscle afferent while monitoring the perception of effort [e.g., 20, 28, 35, 36]. These specific muscle afferents have two important effects on physiological responses to the exercise. First, their discharge ensures adequate cardiorespiratory responses to physical exercise via their role in the exercise pressor reflex, also known as the mechano-metaboreflex [37]. Second, while their inhibitory or excitatory effects on the corticospinal pathway seem to be muscle-dependent [38], recent studies suggest an overall net inhibitory effect on the corticospinal pathway during endurance exercise, thus contributing to the development of neuromuscular fatigue [39]. Moreover, they carry nociceptive signals and thus are involved in the perception of pain and associated discomfort [40]. Since the afferent feedback model considers group III-IV muscle afferents as the signal processed by the brain to generate the perception of effort, pharmacologically blocking muscle afferents should decrease the ratings of perceived ratings of perceived effort. On the other hand, observing stable or increased ratings of perceived effort in the presence of reduced muscle afferent feedback would support a centrally generated perception of effort. Intrathecal and epidural injection of anesthetics or analgesics, such as lidocaine or fentanyl, has traditionally been used to investigate the role of group III-IV muscle afferents and the motor command in cardio-respiratory responses to exercises in both healthy [e.g., 35] and symptomatic participants [e.g., 41] as well as in human performance during endurance exercises [39]. In these studies, participants performed an exercise protocol, usually cycling or isolated knee exercises, with and without intact feedback from group III-IV muscle afferents. While the primary variables of interest were the cardio-respiratory responses to the tasks, these studies often measured the perception of effort as a secondary or tertiary variable. Interestingly, there are conflicting results from these studies. In the presence of pharmacological blockade of muscle afferents, several authors observed a decrease [e.g., 42] in the ratings of perceived effort while others observed no difference [e.g., 36] or an increase [e.g., 43] when compared to a sham or control intervention. This heterogeneity is also found in patients with cardiovascular diseases [e.g., 41, 44]. To the best of our knowledge, only one published article has narratively reviewed the use of pharmacological blockade to explore the neurophysiological mechanisms underlying the perception of effort [28], and a systematic approach has yet to be conducted.

The conflicting findings on the effects of pharmacologically blocking muscle afferents on the perception of effort may be explained by differences in its operational definitions, leading to inconsistencies in the instructions provided to the participants on how to quantify the perception of effort. In his seminal work, Borg defined the perception of effort as how *heavy* and *laborious* a physical task is [45, 46]. However, he also mentioned that this perception results from the integration of various peripheral factors, including the organs of circulation and respiration, the muscles, the skin, and the joints. Since this first definition and associated description provided by Borg, two lines of research have investigated the perception of effort in exercise sciences (see figure 1). A first line of research considers effort as a construct encompassing a mix of exercise-related perceptions [18, 47]. This approach results from the original description proposed by Borg, and later authors supplemented this definition with the notions of fatigue and discomfort: *the subjective intensity of effort, strain, discomfort, and/or fatigue that is experienced during physical exercise* [18]. However, while a lack of familiarization or the presence of fatigue may compromise the dissociation between perceptions of effort and other exercise related perceptions [14, 48–50], experimental data demonstrated that the perception of effort can be dissociated from other exercise-related perceptions, such as pain [51, 52], discomfort [53–55], muscle tension [48, 56] and fatigue [57, 58]. In line with the aforementioned evidence, a second line of research considers effort as a construct dissociated from other perceptions. This approach follows Borg’s original definition conceptualizing the perception of effort as one’s appreciation of the difficulty of a task (how *hard* it is). For instance, Preston and Wegner [59] described the perception of effort as the *feeling of difficulty and labor experienced during exertion*. In 2010, Marcora proposed that the perception of effort is the *conscious sensation of how hard, heavy and strenuous a physical task is* [60]. More recently, Steele defined the perception of effort as the *perception of current task demands relative to the perception of capacity to meet those demands* [16]. In light of the disparate definitions proposed in the literature, it appears crucial to consider the definition used to investigate the perception of effort when interpreting such data. However, to the best of our knowledge, such consideration has not been made in the literature when discussing the signal(s) generating the perception of effort.

**Fig. 1:**
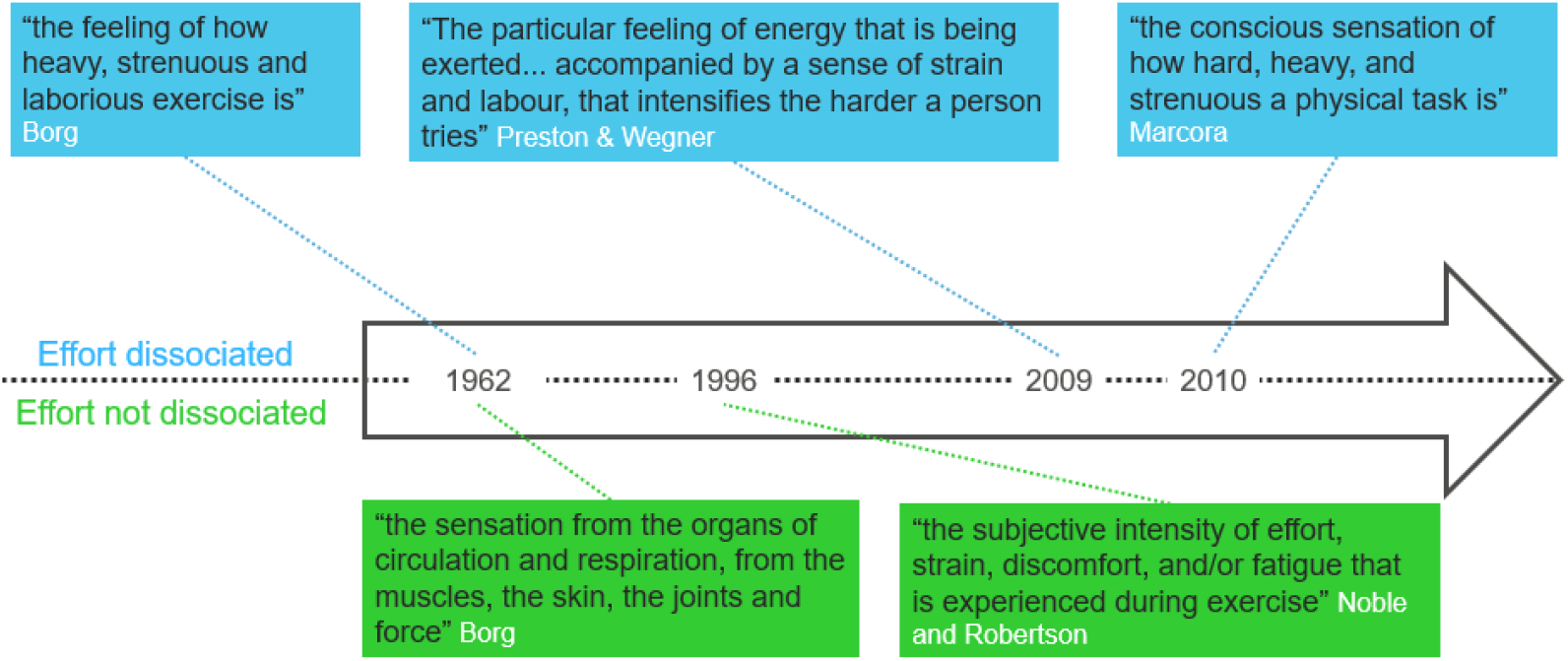
Overview of the two lines of research investigating the perception of effort in exercise sciences and the primary definitions used in the literature. In blue, the perception of effort does not include other exercise-related perceptions and is investigated as a construct dissociated from other perceptions. This approach follows Borg’s original definition conceptualizing the perception of effort as one’s appreciation of the difficulty of a task (how *hard* it is). In green, the perception of effort includes other exercise-related perceptions and is investigated as a construct encompassing a mix of exercise-related perceptions. This ss approach results from the original description proposed by Borg.

In this context, the aim of this systematic review with meta-analysis was to explore the impact of pharmacologically blocking muscle afferents on the perception of effort during physical tasks. To explore whether the inclusion or not of other exercise-related perceptions in the definition of effort influences the quantification of perceived effort, a qualitative analysis was also used to group included studies by their theoretical approach.

## 2. Methods

The present review was conducted and is reported as per the Preferred Reporting Items for Systematic Reviews and Meta-analyses statement (PRISMA; [61]). All methods were pre-specified in a protocol registered on PROSPERO (CRD401913921) prior to the screening process and any deviations from the pre-registered methods are noted throughout.

### 2.1 Search

The following electronic databases were searched to identify studies: MEDLINE, EMBASE, CINAHL, SPORTDiscus, Web of Science and PsycINFO. To ensure the inclusion of recently published articles, the search was conducted on three separate occasions (April 2019 and March 2020 and October 2021). A search strategy was developed for MEDLINE and adapted for each database. The first of the two concepts included was the perception of effort and included the following terms: *perception of effort, sense of effort, perceived exertion, central motor command, central motor drive*. Because the perception of effort has been used by some in the neuroscience literature as an index of the central motor command during exercise, the latter has been included in this concept. The second concept related to the pharmacological blockade and included the following terms: *epidural anesthesia, spinal anesthesia, neural blockade, nerve block, sensory dysesthesia, fentanyl, lidocaine, bupivacaine, muscle afferent, neural feedback, afferent feedback*, and *group III-IV*. It is noteworthy that, despite using the same search strategy in all three instances, PsycINFO returned fewer articles each time the database was scanned (65 against 30 against 27 articles) and that 47 of the 65 original articles were not found in the subsequent searches. Additionally, no limitation to the publication date was set during the search. The complete search strategy for every scanned database is available in the supplementary material S1.

### 2.2 Article inclusion

Eligibility criteria were defined accordingly with the PICOS model. Articles qualified for inclusion if they met the following criteria: 1) *population*: intervention was done on human participants, 2) *intervention*: consisted of a blockade of spinal afferents by epidurally or intrathecally injecting a local anesthetic or analgesics, 3) *comparators*: intervention was compared against a control or placebo condition, 4) *outcome*: the perception of effort was a primary, secondary, or tertiary outcome, and 5) *study design*: repeated-measure designs. Additionally, only articles published in peer-reviewed journals and written in English were retained. We also opted not to include self-paced protocols in the quantitative analyses to allow comparison between studies. This decision is based on the necessity to compare the intensity of the perception of effort when the task demand (e.g., power output) is matched between conditions [16]. The study selection process was done separately by MB and MPDLG, using the online platform Covidence (https://www.covidence.org/home). Conciliation of study selection was done after screening at the title/abstract level and after the full-text screening. Disagreements were settled through discussion and, if necessary, through consultation with BP and JS intervention.

### 2.3 Risks of bias

Risks of bias were appraised with a modified version [62] of the Effective Public Health Practice Project (EPHPP) Quality Assessment Tool for Quantitative Studies [63]. By not applying the selection bias component, this version was adapted to accommodate sport sciences studies where self-referral is common [62]. Further adaptations were made to reflect the need of the present review following the Cochrane Collaboration’s guidelines [64]. The EPHPP does not provide explicit instructions on the assessment of cross-over trials, which was a type of design in the included studies. Appropriately randomized cross-over trials were considered equivalent to randomized controlled trials whereas cross-over trials performed in a counter-balanced order or with inappropriate randomization were considered equivalent to controlled clinical trials. Studies testing participants before and after opioid injection in a pre-post fashion were considered cohort studies. Risks of bias may be different across several outcomes and thus warranted confounders to be appraised specifically for the perception of effort. This is important because authors often consider the perception of effort when interpreting their results [e.g., 65] or use their results to draw conclusions on the regulation of the perception of effort [e.g, 66]. The risk of unblinding in the included studies is high, owing to the side effects the epidural anesthesia may have [e.g., pruritus, dysesthesia; 67]. As such, studies were considered blinded only if they detailed adequate blinding methods, such as the use of a sham injection. Because correctly guessing the interventions may influence the outcomes [68], the reviewers noted whether the participants were asked to guess the order of the interventions when applicable. However, this had no impact on the labelling of the “blinding” component of the EPHPP. The *withdrawals and drop-outs* component was not applied because none was reported in the included studies. This is probably because the included studies were brief, usually spanning over 3 to 5 visits. Finally, all the included studies were either cross-over trial or cohort studies, in which the risk of carry-over is critical. This component was considered when making a judgment for the intervention integrity. As such, the reviewer noted whether the risks of carry-over were minimized by allowing sufficient time in-between experimental visits for the effects of both protocol and the anesthesia to dissipate (wash-out period). Alternatively, risks of carry-over were considered minimal if baseline values between conditions were similar.

### 2.4 Coding process

Included articles were classified into three distinct groups: 1) effort dissociated from other exercise-related perceptions (*effort dissociated* group), 2) effort including other exercise-related perceptions (*effort not-dissociated* group) and 3) not enough information (*unclear* group). A first round attempted to code articles by the reported definition of the perception of effort or the instructions provided to their participants. However, this method was insufficient as none of the studies explicitly reported either of those. Following this, a finer decision-making process was developed by JC, CFB, MB, MPDLG and BP (see figure 2a). During a second round, the reviewers noted whether other exercise-related perceptions(s) were measured (coded as *effort dissociated*) and whether the perception of effort was used interchangeably with other exercise-related perception(s) (coded as *effort not dissociated*). If this information was not available, studies using the perception of effort as an index of the central motor command were coded as *effort dissociated*. Studies that could not be coded on that basis were coded as *unclear*. Articles were separately coded by MB and MPDLG. Decisions were then conciliated, and disagreements settled through discussion with BP. Notably, this process was done separately to the data extraction for meta-analysis (described below). The analyst (JS) was not provided with the final coding until after extraction was complete.

**Fig 2:**
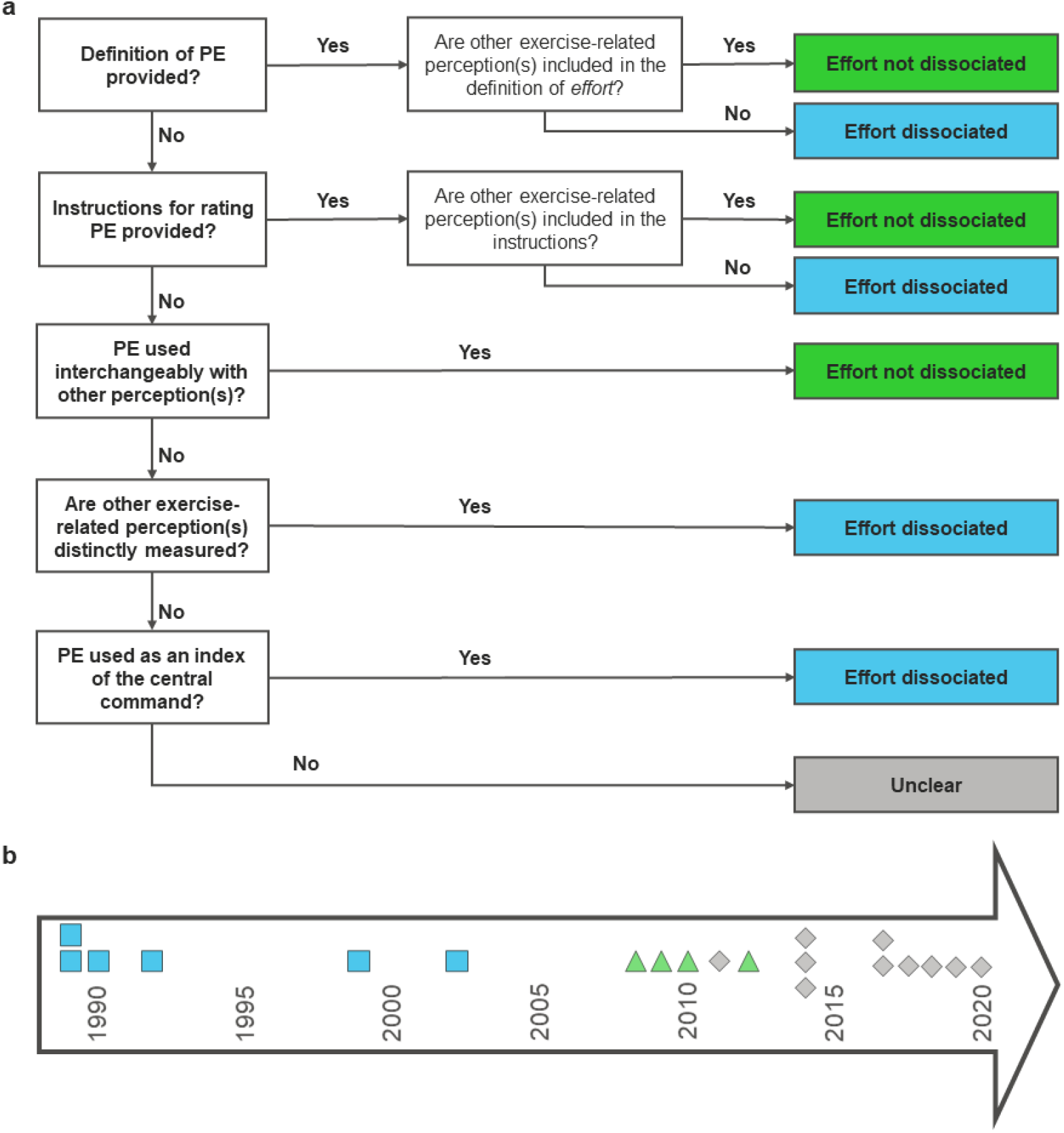
(a) coding process used for the classification of the articles included in the systematic review. (b) changes overtime in the use of the perception of effort construct in the included studies.

### 2.5 Data extraction

Data extraction tables were prepared to map: (a) author and year of publication; (b) the test/exercise conditions used; (c) the method and scale used to capture ratings of perception of effort; (d) the means and standard errors or standard deviations for ratings of perception of effort including both control and intervention conditions; and (e) sample sizes. It is important to note that, due to the multidimensional nature of dyspnea that includes not only respiratory effort but also other physical (chest tightness) and affective (unsatisfied inspiration) components [69, 70], associated ratings were not considered in the present meta-analysis. Of note, we included data for all perception of effort outcomes reported in studies and for all time points for which they were measured; thus, if a study reported multiple perception of effort measurements were captured, these were all included but appropriately coded as either being taken during submaximal tasks or maximal tasks (*i.e*., at task failure). Some studies only reported data graphically or did not report outcomes in a manner conducive to extraction for our analysis, or despite investigating spinal blockade and capturing ratings of perception of effort did not report these outcomes as they were secondary in those studies. In these cases, authors were contacted for sharing either appropriate summary statistics or raw data to facilitate inclusion in our analysis. Three authors were contacted of which one shared the required summary statistics whereas others were either unable or unwilling to share their data. For the studies reporting only graphical data, WebPlotDigitizer (v4.3, Ankit Rohatgi; https://apps.automeris.io/wpd/) was used to extract data for inclusion in our analysis.

### 2.6 Synthesis and analysis

The meta-analysis was performed using the ‘metafor’ [71] package in R (v 4.0.2; R CoreTeam, https://www.r-project.org/). All analysis codes are available on the Open Science Framework website (https://osf.io/cy5n4/). As all studies relied on within-participant design, the standardized mean change using the raw values (SMCR) as described by Becker [72] was calculated between the conditions where the pooled standard deviation from both conditions was used as the denominator for standardization [73]:

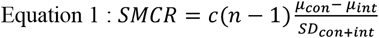

Where *μ_con_* and *μ_int_* are the means for control and intervention conditions respectively, *c* is a bias correction factor [74], and *SD*_(*con+int*)_ is the pooled standard deviation calculated as:

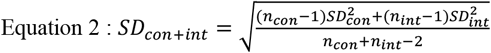

As some studies reported zero variances (*e.g*., some showing ceiling effects), a small constant (six sigma *i.e*., 3*x*10^−7^) was added to all studies prior to calculating effect sizes. The magnitude of standardized effect sizes was interpreted with reference to Cohen [75] thresholds: *trivial* (<0.2), *small* (0.2 to <0.5), *moderate* (0.5 to < 0.8), and *large* (>0.8). Standardized effects were calculated in such a manner that negative effect size values indicated the presence of an intervention effect (*i.e*., a drop in rating of perception of effort), whereas a positive effect size values indicated an effect favoring the control conditions.

As a departure from the pre-registered analysis due to the nested structure of the effect sizes calculated from the studies included (*i.e*., effects nested within groups nested within studies), multilevel mixed effects meta-analyses reflecting these nested random effects in the model were performed. Cluster robust point estimates and precision of those estimates using 95% compatibility (confidence) intervals (CIs) were produced, weighted by inverse sampling variance to account for the within- and between-study variance (tau-squared). Restricted maximal likelihood estimation was used in all models. A main model was produced including all effects sizes. Several exploratory sub-group model analyses were also conducted and detailed in the pre-registration with slight departure here because of the coding process for methods to capture ratings of perception of effort. In addition to a model for all studies, we also produced models for only studies where ratings of perception of effort were captured as *effort dissociated*, *effort not dissociated*, and *unclear* separately. Further, in all models we additionally sub-grouped for tasks as ‘maximal’ (*i.e*., ratings of perception of effort captured at the point of task failure or during a similar maximal task e.g., a maximal voluntary contraction), and ‘submaximal’ (*i.e*., ratings of perception of effort recorded during exercise prior to the point of task failure, or during tasks not necessarily requiring participants to reach task failure e.g., a fixed duration task of absolute demands). This was to separate potential ceiling effects of maximal task conditions. Note, in the preregistration we initially stated we would calculate a pooled effect for all submaximal values reported in any given study. However, to increase the number of effect sizes for analysis, and thus statistical power, we opted to calculate each separately and thus use a multilevel mixed effects meta-analysis model to account for this. We did however also fit the same models as per our pre-registered intention to pool all submaximal values and have included a comparison of these model estimates to those reported here in our supplementary materials which suggests our inferences would not have substantially differed (see https://osf.io/qd6rt/). For each sub-group analysis multilevel models were produced except for the *‘effort not dissociated’* ‘maximal’ conditions where only one effect size was available.

In contrast to our pre-registration, and in light of the heterogeneity and poor reporting of methods to capture ratings of perception of effort, dichotomizing the existence of an effect for the main results was avoided. Therefore, traditional null hypothesis significance testing, which has been extensively critiqued [76, 77], was not employed. Instead, the implications of all results compatible with these data, from the lower limit to the upper limit of the interval estimates, was considered with the greatest interpretive emphasis placed on the point estimate. During revisions however, given some of the apparently null effects identified, we retrospectively refit all models using a Bayesian approach and the ‘brms’ package [78] and produced Bayes Factors using the Savage-Dickey ratio with the ‘bayestestR’ package [79] to compare evidence both for and against a point null of zero difference between conditions. These again did not largely influence our overall inferences (typically weak evidence in favor of the null) are so also included in the supplementary materials with categories of qualitative interpretations as per Kass & Rafferty [80] added to aid interpretation (see https://osf.io/wrkav/).

Risk of small study bias was examined visually through contour-enhanced funnel plots. *Q* and *I*^2^ statistics were also produced and reported [81]. A significant *Q* statistic is typically considered indicative of effects likely not being drawn from a common population. *I*^2^ values indicate the degree of heterogeneity in the effects: 0-40% were not important, 30-60% moderate heterogeneity, 50-90% substantial heterogeneity, and 75-100% considerable heterogeneity [64]. For within participant effects pre-post correlations for measures were rarely reported. Due to the repeated-measure design of included studies, we explored the effect of different correlation coefficients (r = 0.5, 0.7 and 0.9) between pre-post values to test its impact on the results of the models. As overall findings were relatively insensitive to this range, the results for r = 0.7 are reported here. Results for inclusion of the other assumed correlation coefficients are reported in the supplementary material available on OSF (r = 0.5, https://osf.io/gqby6/; r = 0.9, https://osf.io/qbe2n/).

## 3. Results

The search across all databases returned 902 articles. After the removal of 319 duplicates, 583 original articles remained and were screened at the title/abstract level, leaving 80 articles. A total of 20 articles were retained. The screening process is shown in figure 3. None of the included articles defined the perception of effort nor provided instructions for rating it. Therefore, included studies could not be coded on that basis and instead, other cues have been used (see figure 2a). Based upon this, among the 20 articles, only 6 could be classified in the *effort dissociated* group, while 3 were placed in the *effort not dissociated* group and 11 in the *unclear* group. A timeline of the included studies is available in figure 2b. Three studies, 1 in the *effort not dissociated* group and 2 in the *unclear* group used a self-paced protocol and could thus not be added to quantitative analyses. Data was not available for quantitative synthesis in 1 study belonging to the *unclear* group, providing a sample size on which quantitative synthesis was based of n = 164. Full details of all studies included in quantitative synthesis can be seen in the data extraction table (https://osf.io/ku3w7/).

**Fig 3:**
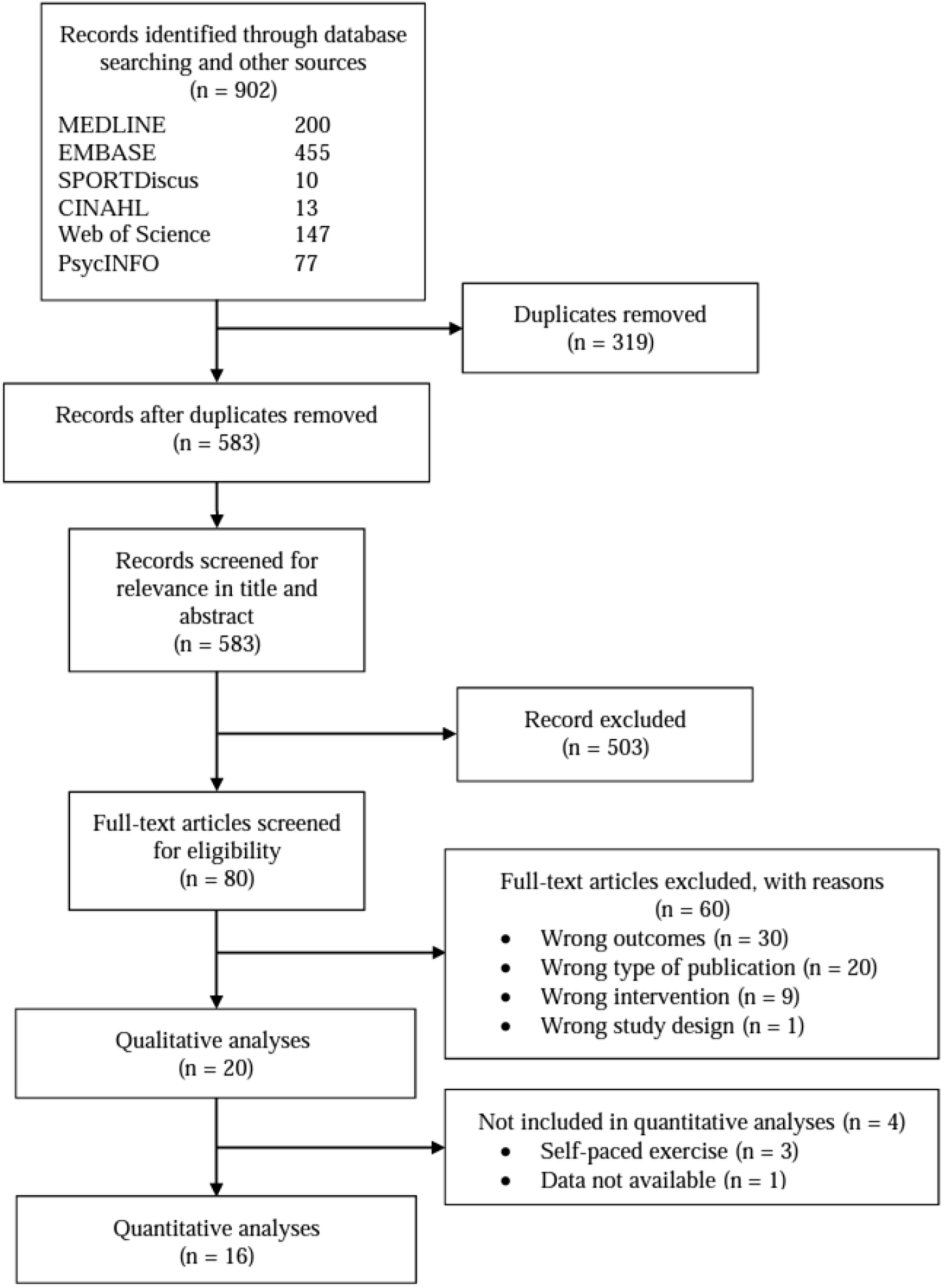
Flow diagram of systematic review inclusion/exclusion adapted from the Preferred Reporting Items for Systematic Reviews and Meta-Analyses.

### 3.1 Risks of bias

A modified version of the EPHPP to accommodate sport sciences studies has been used to assess the risk of bias of individual studies [62, 82]. Studies labelled “strong” are considered to have guarded themselves well against biases. Conversely, studies labelled “weak” are considered to have a high risk of bias. None of the studies were labelled “strong”, whereas 5 were “moderate” and 13 “weak”. Most studies used a randomized or counter-balanced within-subject design, but 7 studies were classified as cohort studies (pre-test, post-test). Almost half (n = 7) of the studies were deemed to have strongly controlled for known confounders of the perception of effort, but 6 of them were labelled “moderate”, with the remaining 5 being considered “weak”. None of the studies described how the assessor was blinded except for one [83]. In contrast, 5 studies blinded their participants with a placebo injection. The remaining 13 studies compared the epidural anaesthesia with a “no-intervention” control condition and the participants could therefore not be considered blind. Furthermore, none of the included studies reported asking the participants to guess the order of the intervention after completion of the protocols. All studies used either the Borg’s scale (RPE 6-20) or the CR10 scale, which are known psychophysical scales in the context of physical exercise. However, several (n = 9) articles reported to have used a modified version of the CR10 scale without providing any information on the modifications performed. The validity and reliability of these modifications were therefore unclear. The table 1 shows the risk of bias within studies for every included articles. An audit trail is available in the supplementary material 2.

**Table 1:**
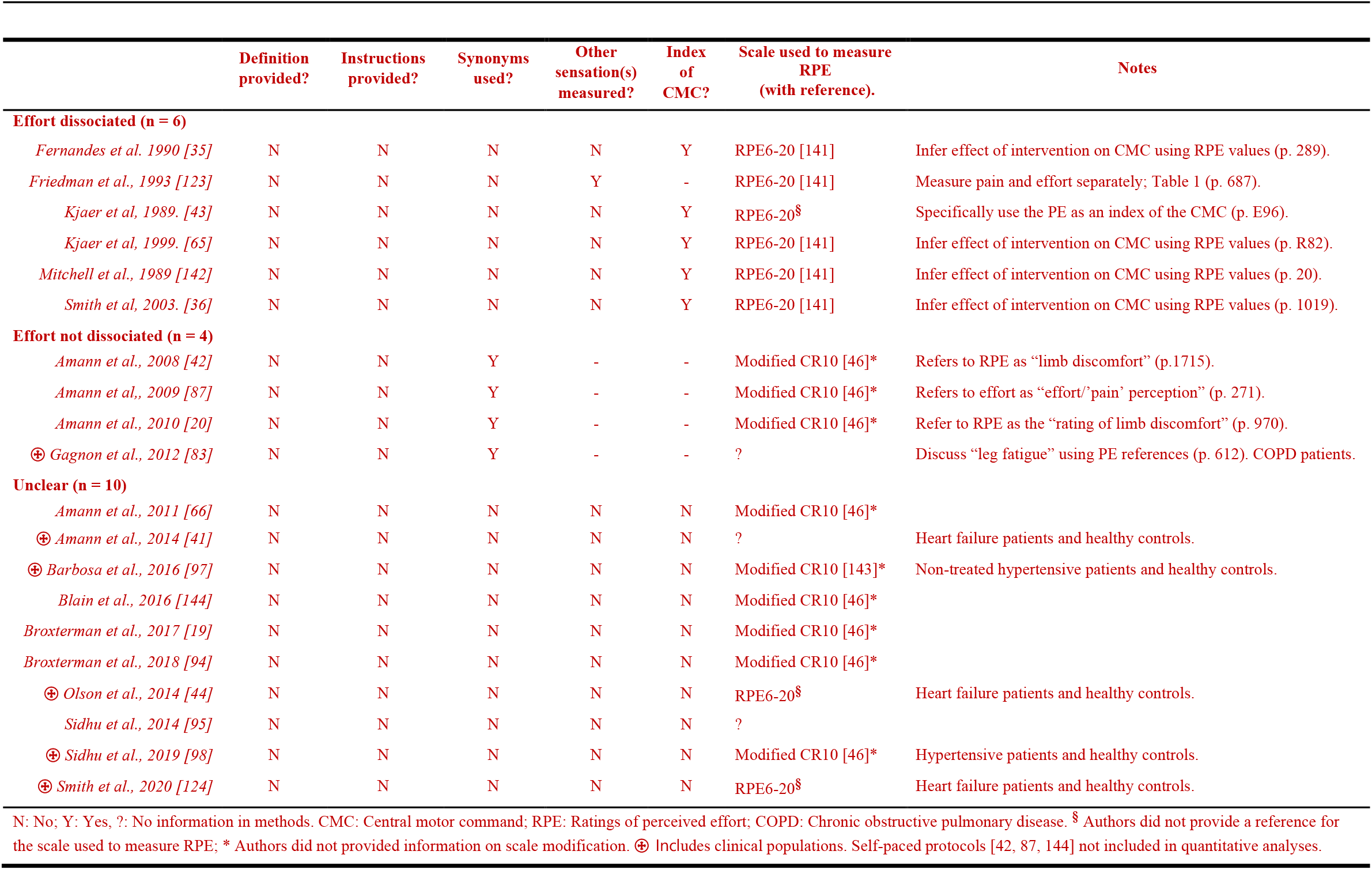
Coding criteria for individual studies

**Table 2:**
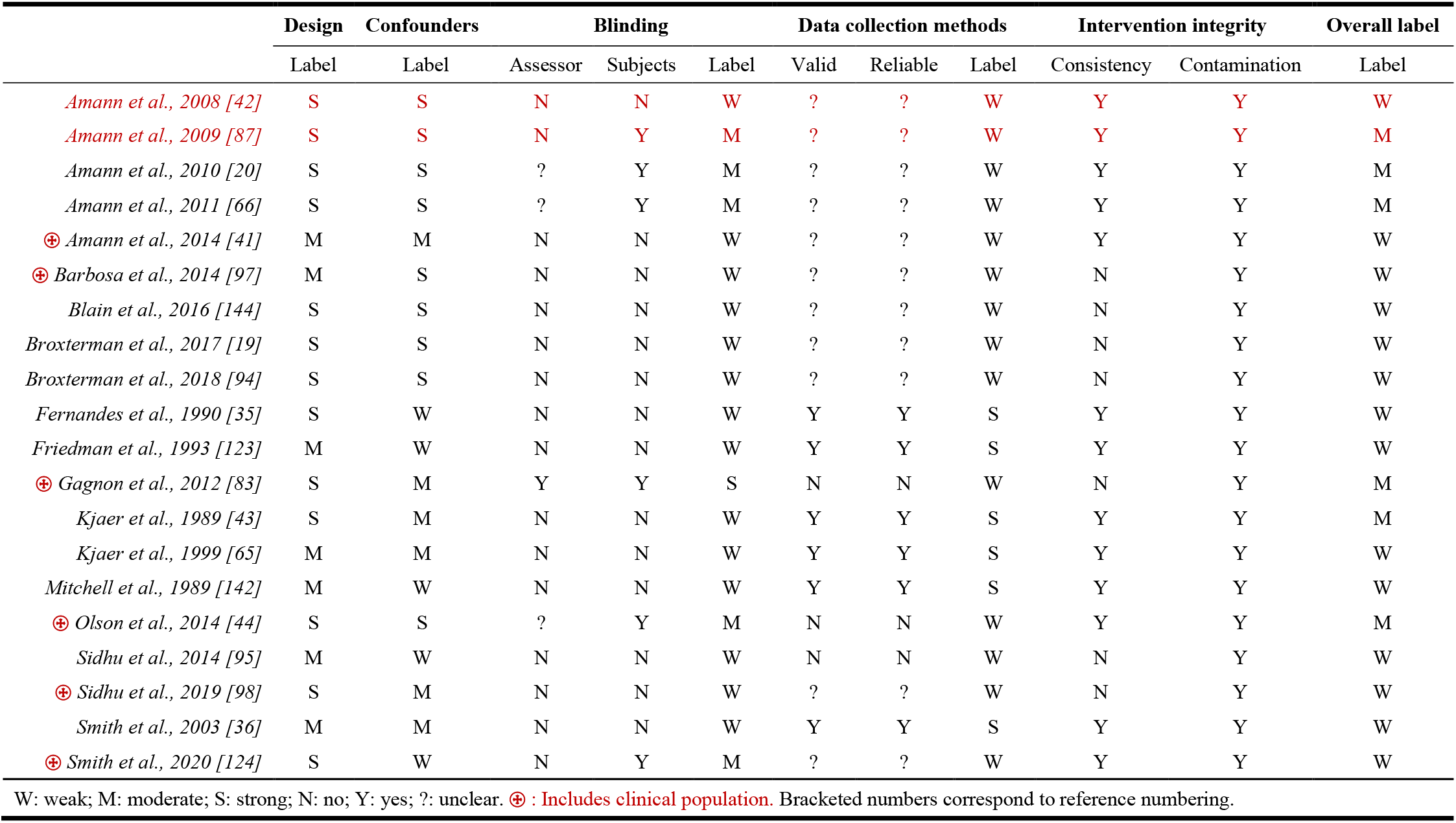
Risks of bias within included studies

### 3.2 Meta-Analysis

The quantitative analysis could be performed on only 16 studies, the remaining two providing insufficient information. The main model including all combined effects sizes (*k* = 49 across 16 clusters [median = 2, range = 1 to 8 effects per cluster]) revealed a negative trivial point estimate with precision ranging from negative small to positive trivial effects for the interval estimate (−0.05 [95% CI = −0.28 to 0.18]), yet with moderate to substantial heterogeneity (*Q*_(48)_ = 127.22, p < 0.0001, *I*^2^_between_study_ = 77%, *I*^2^_between_group_ 0%, *I*^2^_within_group_ = 0%). When considering only submaximal conditions the model (*k* = 39 across 13 clusters [median = 2, range = 1 to 8 effects per cluster]) revealed a point estimate close to zero with precision ranging from negative small to positive small effects for the interval estimate (0.05 [95% CI = −0.32 to 0.42]), with substantial heterogeneity (*Q*_(38)_ = 103.00, p < 0.0001, *I*^2^_between_study_ = 77%, *I*^2^_between_group_ = 0%, *I*^2^_within_group_ = 0%). Considering only maximal conditions the model (*k* = 10 across 9 clusters [median = 1, range = 1 to 2 effects per cluster]) revealed a negative small point estimate with precision ranging from negative moderate to positive trivial effects for the interval estimate (−0.17 [95% CI = −0.47 to 0.13]), with moderate to substantial heterogeneity (*Q*_(9)_ = 22.92, p = 0.0064, *I*^2^_between_study_ = 0%, *I*^2^_between_group_ = 30%, *I*^2^_within_group_ = 30%). Figure 4 presents the funnel plot for all studies with color coding by method for capturing rating of perception of effort. Figures 5 and 6 present all effect sizes for the submaximal models and the overall model estimates for all coding and combined.

**Fig 4:**
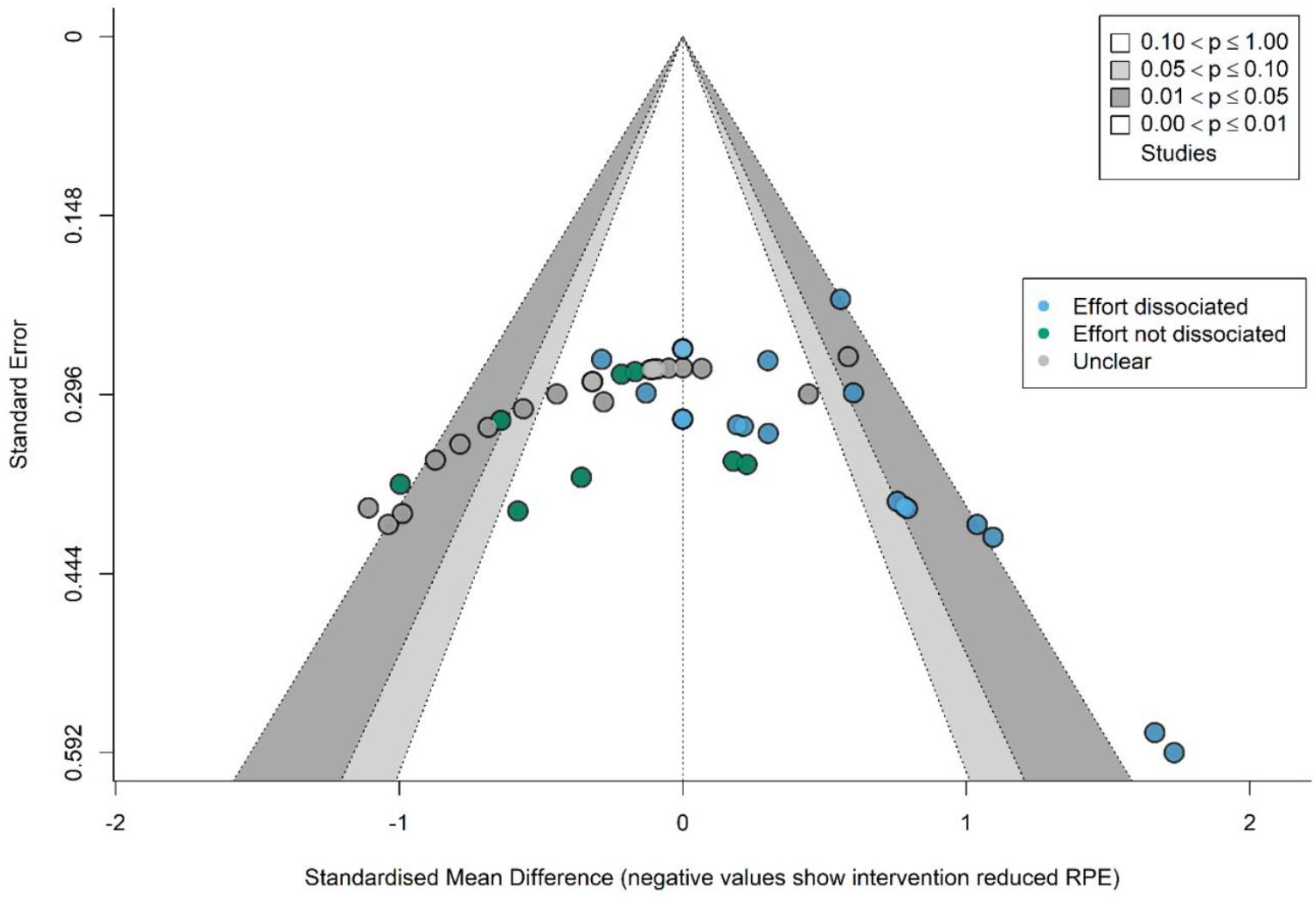
Contour enhanced funnel plot for all effects (*i.e*., rating of perception of effort as *effort dissociated, effort not-dissociated, unclear* color coded; see key).

**Fig 5:**
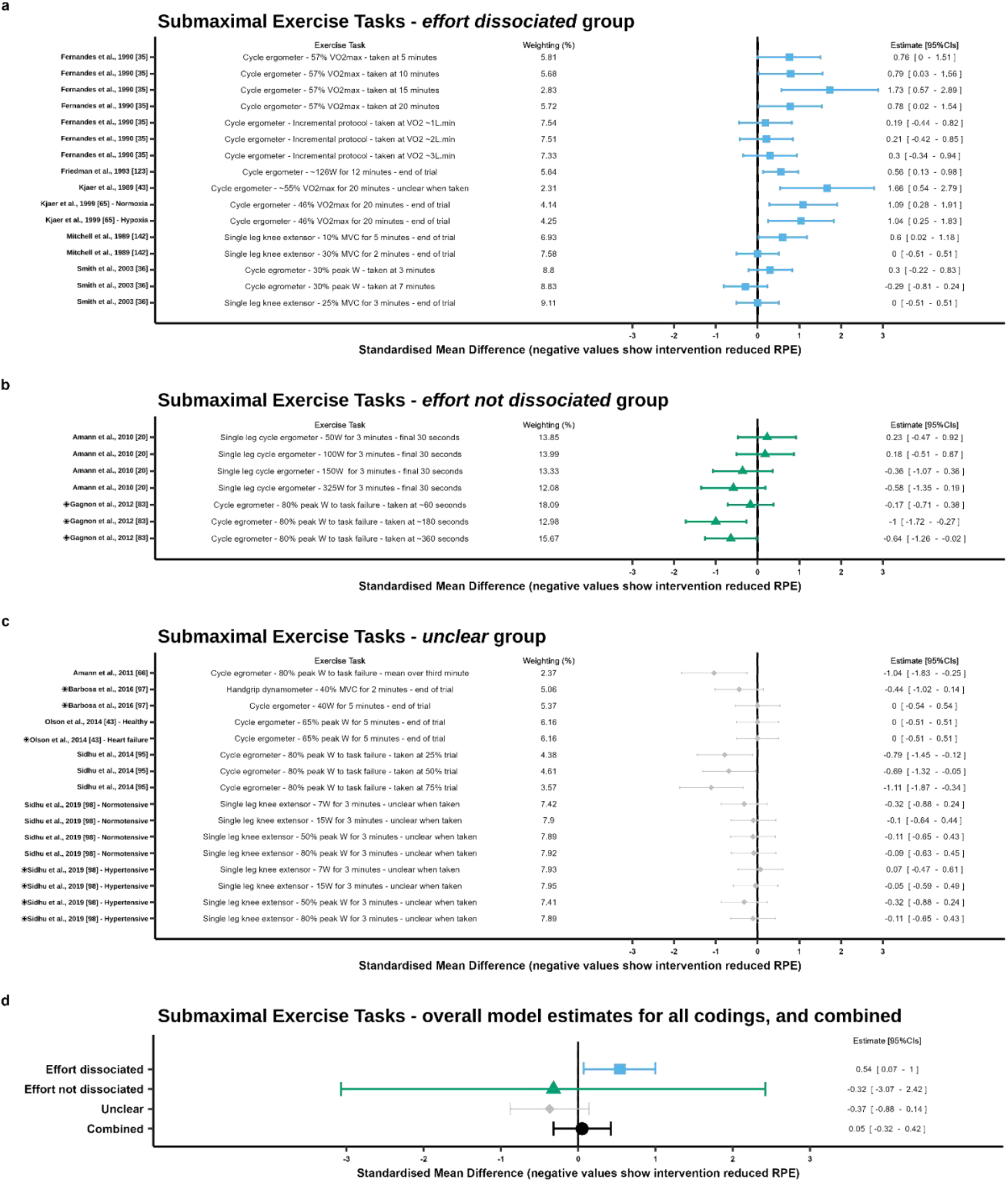
Forest plots for the effect of an epidural anesthesia on the perception of effort at submaximal exercise demands (comparison epidural vs. placebo or no intervention). (a) effect sizes for the *effort dissociated* group. (b) effect sizes for the *effort not dissociated* group. (c) effect sizes for the *unclear* group. (d) overall effect sizes for all coding and combined. Standardized mean differences with 95% confidence intervals are shown.

**Fig 6:**
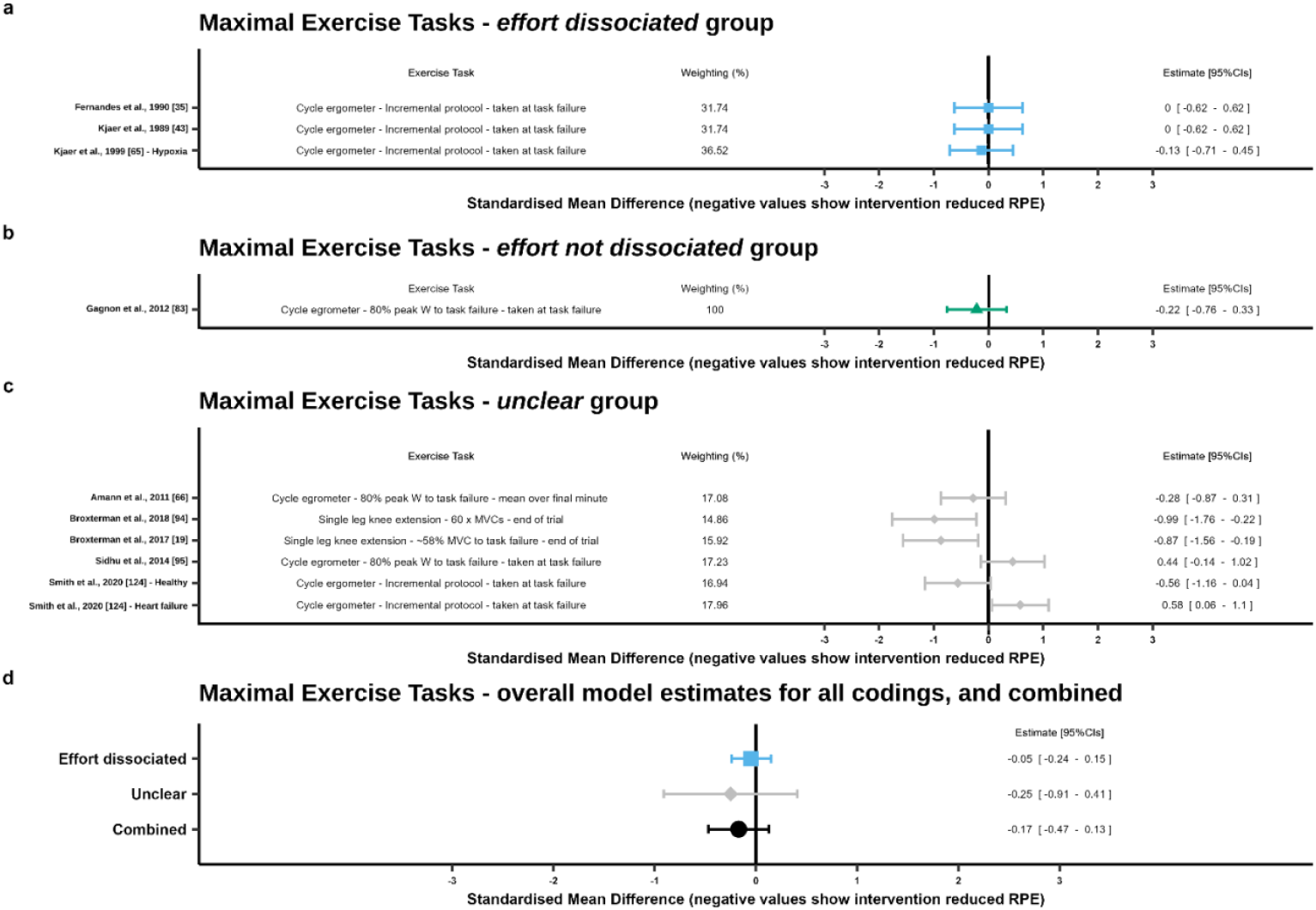
Forest plot for the effect of an epidural anesthesia on the perception of effort at maximal demands (comparison epidural vs. placebo or no intervention). (a) effect sizes for the *effort dissociated* group. (b) effect sizes for the *effort not dissociated* group. (c) effect sizes for the *unclear* group. (d) overall effect sizes for all coding and combined. Standardized mean differences with 95% confidence intervals are shown.

#### Ratings of perception of effort as ‘effort dissociated’

The subgroup model where the method of rating of perception of effort was coded as *effort* (*k* = 19 across 6 clusters [median = 2.5, range = 1 to 8 effects per cluster]) revealed an overall small point estimate with precision ranging from positive trivial to positive moderate effects for the interval estimate (0.39 [95% CI = 0.13 to 0.64]), with relative homogeneity (*Q*_(18)_ = 36.96, p = 0.0053, *I*^2^_between_study_ = 77%, *I*^2^_between_group_ = 0%, *I*^2^_within_group_ = 49%). When considering only submaximal conditions the model (*k* = 16 across 6 clusters [median = 2, range = 1 to 7 effects per cluster]) revealed a moderate point estimate with precision ranging from positive trivial to positive large effects for the interval estimate (0.54 [95% CI = 0.07 to 1.0]), with moderate to substantial heterogeneity (*Q*_(15)_ = 31.44, p = 0.0077, *I*^2^_between_study_ = 51%, *I*^2^_between_group_ = 0%, *I*^2^_within_group_ = 8%). When considering only maximal conditions the model (*k* = 3 across 3 clusters [1 effect per cluster]) revealed a negative trivial point estimate with precision ranging from negative small to positive trivial effects for the interval estimate (−0.05 [95% CI = −0.24 to 0.15]), with relative homogeneity (Q_(2)_ = 0.12, p = 0.9412, *I*^2^_between_study_ = 0%, *I*^2^_between_group_ 0%, *I*^2^_within_group_ = 0%).

#### Ratings of perception of effort as ‘effort not dissociated’

The subgroup model where the method of rating of perception of effort capture was coded as *effort not dissociated* (*k* = 8 across 2 clusters [4 effects per cluster]) revealed an overall negative small point estimate with poor precision ranging from negative large to positive large effects for the interval estimate (−0.29 [95% CI = −2.39 to 1.8]), with relative homogeneity (*Q*_(7)_ = 9.62, p = 0.2113, *I*^2^_between_study_ = 7%, *I*^2^_between_group_ = 7%, *I*^2^_within_group_ = 14%). When considering only submaximal conditions the model (*k* = 7 across 2 clusters [median = 3.5, range = 3 to 4 effects per cluster]) revealed a negative small point estimate poor precision ranging from negative large to positive large effects for the interval estimate (−0.32 [95% CI = −3.07 to 2.42]), with moderate heterogeneity (*Q*_(6)_ = 9.51, p = 0.1467, *I*^2^_between_study_ = 12%, *I*^2^_between_group_ = 12%, *I*^2^_within_group_ = 20%). When considering only maximal conditions only a single effect met these conditions which revealed a negative small point estimate with precision ranging from negative moderate to positive small effects for the interval estimate (−0.22 [95% CI = −0.76 to 0.33]).

#### Ratings of perception of effort as ‘unclear’

The subgroup model where the method of rating of perception of effort capture was coded as *unclear* (*k* = 22 across 8 clusters [median = 2, range = 1 to 8 effects per cluster]) revealed an overall negative small point estimate with precision ranging from negative moderate to negative trivial effects for the interval estimate (−0.27 [95% CI = −0.50 to −0.04]), with moderate heterogeneity (*Q*_(21)_ = 42.14, p = 0.004, *I*^2^_between_study_ = 6%, *I*^2^_between_group_ = 0%, *I*^2^_within_group_ = 45%). When considering only submaximal conditions the model (*k* = 16 across 5 clusters [median = 2, range = 1 to 8 effects per cluster]) revealed a negative small point estimate with precision ranging from negative large to positive trivial effects for the interval estimate (−0.37 [95% CI = −0.88 to 0.14]), with moderate to substantial heterogeneity (*Q*_(15)_ = 19.27, p = 0.2020, *I*^2^_between_study_ = 59%, *I*^2^_between_group_ = 0%, *I*^2^_within_group_ = 0%). When considering only maximal conditions the model (*k* = 6 across 5 clusters [median = 1, range = 1 to 2 effects per cluster]) revealed a negative small point estimate with poor precision ranging from negative large to positive moderate effects for the interval estimate (−0.25 [95% CI = −0.91 to 0.41]), with considerable heterogeneity (*Q*_(5)_ = 22.47, p = 0.004, *I*^2^_between_study_ = 0%, *I*^2^_between_group_ = 39%, *I*^2^_within_group_ = 39%).

## 4. Discussion

This systematic review with meta-analysis was conducted to investigate the effect of pharmacologically blocking muscle afferents on the perception of effort during physical exercise. We also sought to explore the differences in reported ratings of perceived effort according to the approach used to investigate this construct. Twenty articles were coded as *effort dissociated* (n = 6), *effort not dissociated* (n = 4) or *unclear* (n = 10) according to whether the authors included other exercise-related perceptions in the investigation of the perception of effort. Considering all subgroups combined, a trivial negative point estimate for ratings of perceived effort reduction following pharmacological blockade of muscle afferents was observed, with interval estimate ranging from negative small to positive trivial effects. The magnitude of the interval estimate crossing zero as well as the trivial negative point of estimate (0.05) suggests that regardless of the theoretical approach used to investigate the perception of effort, pharmacologically blocking muscle afferents feedback does not reduce the perception of effort during physical exercise.

The subgroup analysis also revealed a clear influence of the theoretical approach used to investigate the perception of effort (i.e., as a construct dissociated or not from other exercise-related perceptions). To the best of our knowledge, this systematic review is the first to highlight an influence of the theoretical approach used to investigate this perception, and strongly suggests that future experimental studies should carefully report the instructions provided to the participants for rating perception of effort.

### 4.1 Effort dissociated subgroup

Only considering the *effort dissociated* subgroup, pooling 6 studies yielded a small positive point estimate with a positive confidence interval suggesting that in the presence of impaired group III-IV muscle afferents, participants may report higher ratings of perceived effort. Participants working at the same absolute demands (*i.e*., same external workload in both conditions) reported higher perception of effort with epidural anesthesia [35, 36, 43]. However, when working at similar relative workloads (i.e., taking into consideration any muscle strength reduction following injection of local anesthetic [42]), the same participants reported similar perception of effort. Likewise, ratings of perceived effort were also similar at a given oxygen uptake during graded exercises [35, 43]. When only considering submaximal tasks, the *effort dissociated* subgroup also showed moderate heterogeneity. Differences across studies could be explained by differences in experimental designs [84], as results differed with the calibration of exercise demands according to absolute or relative workloads. It is also important to note that the increased perception of effort in these studies is likely due to the use of lidocaine and/or bupivacaine to block III-IV muscle afferent feedback. Indeed, contrary to fentanyl which acts more specifically on sensory transmission [85], lidocaine and bupivacaine is known to affect sodium and potassium channels [86] and reduce force production capacity [87] thereby requiring the participants to increase their motor command to maintain the same absolute workload. According to the corollary discharge model, this increased motor command increases the magnitude of the associated corollary discharge, which in turn increases the perception of effort [24]. This subgroup analysis reveals that when effort is investigated as dissociated from other exercise-related perceptions, pharmacological blockade of muscle afferents does not reduce perception of effort. Therefore, as perception of effort is not reduced, muscle afferent feedback cannot be considered as a sensory signal processed by the brain to generate the perception of effort. This result reinforces the potential of using the perception of effort intensity as a psychophysiological index of the motor command [24, 27, 88–90], as traditionally performed in the neuroscience, cardiovascular physiology and kinesthesia literatures [24, 25, 27, 91].

### 4.2 Effort not-dissociated subgroup

Only considering the *effort not dissociated* subgroup, pooling 2 studies yielded an overall small negative point estimate. This negative point estimate was associated with an important imprecision based upon the confidence interval range likely due to the low number of studies and small cluster sample correction for robust estimates. Both studies observed lower ratings of perceived effort in epidurally anaesthetized participants [20, 83]. Interestingly, Amann et al. [20] observed lower ratings of perceived effort only at higher cycling power output (*i.e*., 80% peak power output, 325 ± 19 W). According to the Oxford Dictionary, discomfort can be defined as a slight pain and something that makes a person feel physically uncomfortable. Because of its relation to the concept of pain, the inclusion of discomfort in the definition of the perception of effort may bias the ratings of perceived effort whenever there is a change in the perception of pain [14, 92]. Although there seems to exist wide interindividual variability in pain threshold during cycle ergometry, muscle pain is known to increase with increased exercise demands during this task [93]. Attending to these perceptions when measuring the perception of effort may attenuate the perceptual differences between conditions during lower cycling demands where discomfort is already low and thus, any difference with epidural anesthesia would likely be minimal. Conversely, when working at 80% peak power output, participants reported substantially lower ratings of perceived effort with epidural anesthesia, probably because they felt less discomfort and/or pain than they normally would have. Pain and associated unpleasant sensations are transmitted through group III-IV fibres [40], and thus are attenuated with epidural anesthesia. Amann et al., [20] and Gagnon et al. [83] both observed lower ratings of perceived effort, suggesting that the reduction in muscle pain and associated discomfort may have biased the ratings of perceived effort when other exercise-related perceptions were included in the definition of effort. A study employing a self-paced protocol that could not be added to this meta-analysis found similar results [42]. Participants performed a 5-km cycling time trial with and without lidocaine with a mean power output similar to that of Amann et al. [20]. The authors however found a decrease in ratings of perceived effort of nearly 2 unit-points on the CR10 scale (CTRL: 8.4 ± 0.4 SEM; Lidocaine: 6.8 ± 0.4 SEM). Given the high-power output, the fact that feedback from group III-IV is known to be the signal processed by the brain to generate muscle pain [40], and that the authors reported investigating “limb discomfort”, the lower values likely reflect a decreased pain and discomfort when cycling with lidocaine. As such, the inclusion of discomfort in the definition of effort would likely result in a decrease in reported ratings of perceived effort. Interestingly, a similar protocol, with the only difference of using fentanyl instead of lidocaine, observed an increase in ratings of perceived effort at the completion of the 5 km time-trial [66]. However, the authors also observed an excessive development of fatigue resulting in an increase in central motor drive, known to exacerbate the perception of effort [4, 24, 27, 30]. Moreover, when an effect of the epidural anesthesia is detected (e.g., 13% reduction in “limb discomfort”[20]), the magnitude of its effect is small. Even when the theoretical approach encompasses several exercise-related perceptions, the contribution of group III-IV muscle afferents appears to be limited as previously suggested [92].

### 4.3 Unclear subgroup

Only considering the *unclear* subgroup, pooling 8 articles yielded an overall small negative point estimate with a negative confidence interval. Among those articles, 2 observed lower ratings of perceived effort at task failure when exercising with impaired muscle afferents with similar integrated forces between conditions [19, 94]. Interestingly, 2 other studies observed similar ratings of perceived effort at task failure. Amann et al. [66] found a 27% decrease in perception of effort intensity with epidural anesthesia at the 3-min mark (average time, placebo: 8.7 ± 0.3, fentanyl: 6.8 ± 0.3), but not at exhaustion in trained athletes cycling at 80% of their peak power output. Similarly, Sidhu et al. [95] observed a decrease in perception of effort intensity with epidural anesthesia only at 25% of endurance time during a similar exercise protocol. It must be noted that a 1-unit difference on the CR10 was consistently maintained throughout the protocol until exhaustion where values were nearly identical (100% ET, 9.9 ± .01 vs 10.0 ± 0). It is indicative of a tendency for ratings of perceived effort to be lower in the epidural anesthesia condition, albeit not reaching statistical significance. Similar values at the end of the endurance time would also be consistent with previous studies suggesting that the perception of effort attain near maximal values at exhaustion [e.g., 96]. Three other studies using fentanyl in hypertensive and heart failure patients also did not find different ratings of perceived effort compared to a sham or control condition [44, 97, 98]. Exercise demands, determined with cycling power output, was extremely low in one of the studies (40 W; [97]). Because participants reported similar ratings of perceived effort when also considering discomfort at lower intensities [20] and that fentanyl does not lead to loss in muscle strength [87], it is not possible to determine which was the cause of the lack of differences. However, when exercising at higher intensities (*e.g*., 65% −80% peak power output), healthy controls and heart failure participants reported similar ratings of perceived effort during both conditions [44]. Furthermore, healthy and heart failure participants did not differ in ratings of perceived effort despite large differences in external workload [44], further suggesting that it is the relative and not the absolute workload determining the perception of effort.

Although the involvement of the central motor drive in the perception of effort seems widely accepted [24, 42], there is still confusion about the role of group III-IV muscle afferents as a signal processed by the brain to generate the perception of effort [14, 29]. In fact, physiologists have still not reached a consensus after more than 150 years of debate [99–102]. Central projections of group III-IV muscle afferents to several spinal and supra-spinal sites, including sensory cortices, anatomically support the afferent feedback model [103, 104]. This model also finds experimental evidence from studies involving epidural anesthesia [20, 83]. However, as mentioned in the introduction, the inclusion of other exercise-related perceptions likely biased the ratings of perceived effort measured. Furthermore, if these muscle afferents constitute a centrally processed signal generating the perception of effort, stimulation of group III-IV muscle afferents would generate a sense of effort even in the absence of central motor drive. However, injections of physiological concentrations of metabolites known to stimulate those muscle afferents do not generate perception of effort at rest [105, 106]. This manipulation however elicits sensations related to discomfort (*e.g*., itch, tingling) and pain. It appears that the presence of the motor command, and therefore the voluntary engagement of the participant in the task, is crucial for experiencing the perception of effort.

### 4.4 Other (neuro-)physiological signals potentially processed by the brain to generate the perception of effort

As shown in figure 1, Borg’s original description [45, 46] suggest that the perception of effort may also be generated by the brain processing of several peripheral inputs, including organs of circulation and respiration, skin and joints. Regarding the organs of circulation and respiration, there is evidence that heart and lung transplant recipients (i.e., denervated organs) may perceive effort normally [107, 108]. Moreover, administration of β-blockers prior to exercise does not reduce the perception of effort despite a decrease in heart rate [109]. Therefore, it seems that afferent feedback from the heart and the lungs is not processed by the brain to generate the perception of effort. Regarding skin and joint feedback, evidence from the kinesthesia literature suggests that this specific feedback is involved in the perceptions of force and movement rather than the perception of effort [26]. Finally, other authors have proposed that hormones and cytokines may mediate the perception of effort [e.g., 110, 111]. It is important to remind that the perception of effort is instantaneously experienced during the voluntary engagement in a physical task and immediately disappears when an individual disengages from it. Given the relatively slow-acting nature of hormones and cytokines (e.g., secretion, circulation to the brain, crossing of the blood-brain barrier), it is unlikely that their signal is processed by the brain to generate the perception of effort. However, it is plausible that hormones and cytokines may indirectly interact with the perception of effort, for example by altering the neuronal processing of the signal generating the perception of effort.

There is extensive support that the neurocognitive processing of the corollary discharge generates the perception of effort. Stimulation of muscle afferents, in the absence of motor command, generates various perceptions (e.g., pain, movement, force), but not effort [105, 106, 112, 113]. Moreover, significant correlations between the ratings of perceived effort and the amplitude of movement-related cortical potential (MRCP), an index of the motor command, have previously been observed [24, 27, 88, 114, 115]. For example, a reduction in force production capacity is associated with an increased MRCP amplitude and an increased perception of effort intensity to maintain the same absolute force [24]. When caffeine is ingested, perception of effort intensity to maintain the same absolute force is reduced in association with a decreased MRCP amplitude [27]. This positive effect of caffeine on the perception of effort is most likely due to the increased excitability of the corticospinal pathway induced by its ingestion [116–120], leading to a lower motor command required to activate the working muscles as revealed by the decreased MRCP amplitude. Second, support in favor of the role of the corollary discharge as an internal signal generating the perception of effort can be found in various neuroscience or psychophysiological studies. For example, Zenon et al. [121] demonstrated that disrupting the supplementary motor area via continuous theta burst transcranial magnetic stimulation decreases perception of effort. Other studies demonstrated a close relationship between perception of effort and physiological variables known to be strongly influenced by the motor command, such as the respiratory frequency [122] or the electromyographic signal [90].

### 4.5 Strengths and limitations

One strength of this systematic review is that our search was not restrained by publication date, despite the articles spanning three decades. Rather, the shift from muscle weakness-inducing lidocaine and bupivacaine to the highly selective *μ*-opioid receptor agonist fentanyl demonstrated the importance of the magnitude of the central motor drive in generating the perception of effort. In the presence of muscle weakness (induced by some opioids binding to spinal motoneurons), exercisers must increase the magnitude of their motor command to maintain the same absolute level of performance [32]. This was observed in older studies via the higher ratings of perceived effort reported when participants were working at similar external workload [e.g., 43, 123]. Furthermore, a shift in the usage of the perception of effort, from *effort dissociated* to *effort not dissociated* and *unclear* is observed (figure 2b). Another strength of this meta-analysis is that we opted to pool the data per time-point, instead of averaging the ratings of perceived effort within studies. This allows us to avoid any potential ceiling effects from maximal conditions, where ratings of perceived effort are expected to attain near maximal values. Moreover, this approach offered a view of the effect of pharmacologically blocking group III/IV muscle afferents at different exercise demands, as several authors employed incremental protocols. This proved particularly useful to interpret data from the *effort not dissociated* subgroup, as discomfort and pain, often included in the definition of effort, are more predominant at higher workloads.

One limitation of this systematic review is that none of the included studies explicitly provided the definition of the perception of effort, or the instructions given to the participants to report their ratings of perceived effort. Therefore, to overcome this limitation, a unique coding was created to be able to classify the included articles based on cues leading to the assumption or not of the inclusion of other exercise-related perceptions in the definition of effort. While some may argue that such approach opens the door for interpretation, we would like to emphasize that the coding process was clear and objective, and could be reproduced by other research groups, as presented in figure 2a. We are therefore confident that our coding process was successful at separating studies according to the theoretical approach used by the researcher (dissociated perception or not). Importantly, this limitation strongly emphasizes the need for better reporting of the definition of the perception of effort and associated instructions provided to the participants directly in the manuscript, or in supplementary materials. Indeed, in some manuscript, information concerning the perception of effort solely appeared in the results without any further detail [e.g., 124]. Another important point is that 9 out of 20 studies included in the qualitative analysis (~ 50%) reported the use of modified scales to measure the perception of effort, without providing these modifications. Explicitly reporting definition and instructions, as well as the psychophysical scale used, is fundamental for study reproducibility. Such rigor should decrease the heterogeneity in the results, regardless of the theoretical approach chosen by the research groups.

Because of the contrasting effects of group III/IV muscle afferents on different spinal and supra-spinal networks [38, 39], it can be difficult to make clear conclusion on their role in the regulation of the motor command and the perception of effort. Importantly, we would like to emphasize that our systematic review and meta-analysis investigated the role of group III-IV muscle afferent as a potential signal processed by the brain to <u>generate</u> the perception of effort (i.e., required for perception of effort to be experienced). According to this, blocking this signal (i.e., generator) should abolish, or at least considerably reduce the perception of effort. By combining our results with existing evidence that stimulation of these afferents in the absence of motor command does not generate a perception of effort [105, 106], it seems clear that feedback from group III-IV muscle afferents is not processed by the brain to generate this perception. Stimulation of these muscle afferents generates other perceptions, and particularly pain [40, 105]. To reinforce this point, we could draw a comparison with the perception of movement. Movement perception is well known to be generated by the brain processing of both the copy of the motor command and muscle afferent feedback [26, 125, 126]. In the absence of motor command, stimulation of muscle afferents via tendon vibration creates an illusion of movement [e.g., 112, 113]. As previously mentioned, stimulation of muscle afferents in the absence of motor command, according to the existing literature, does not create an illusion of effort when effort is defined as a perception dissociated from other exercise related perceptions. While our results and the literature exclude the possibility that feedback from group III-IV muscle afferents is a signal processed by the brain to generate the perception of effort, we would like to emphasize that these afferents can play an indirect role in the regulation of this perception. Indeed, as previously mentioned, feedback from group III-IV muscle afferent interacts with the regulation of cardiorespiratory responses to the exercise and the regulation of the motor command sent to the working muscles [37, 39]. Consequently, it is important for future research to further investigate how this feedback modulate (as opposed to “generate”) the perception of effort in healthy and clinical populations. An interesting approach could consist of increasing feedback from group III-IV muscle afferents concomitantly of the presence of the perception of effort. This is possible for example with the use of intramuscular metabolites or saline injection [e.g., 127, 128, 129] or cuff-induced muscle ischemia [e.g., 130, 131, 132]. Due to the overall inhibitory effect of group III/IV muscle afferents on the corticospinal pathway and the generation of the motor command, thus contributing to the development of neuromuscular fatigue [38, 39], we would expect an increase in the perception of effort.

Finally, recent literature questions the inclusion of *heaviness* in the definition of effort [16, 133]. This debate is in part related to the link between the word *heavy* and the weight of an object to lift. However, the word *heavy*, as defined by the Oxford dictionary, refers to “the quality of having great weight” (i.e., weight of an object) as well as “a state of being greater in amount in force, or intensity than usual” (i.e., perceived task demand). Because this debate is fairly recent and the word *heaviness* is included in Borg’s original definition, separating *heaviness* from effort would have been too restrictive. Thus, this was not considered for the present meta-analysis. Considering this interesting debate on the influence of the inclusion of the word “heaviness” in the perception of effort, we propose two avenues. First, future research should investigate the effect of including or not the word *heaviness* or *heavy* in the rating of perceived effort in various exercise types, including lifting weight (e.g., resistance exercise) or not (e.g., endurance exercise). Second, future research using current definitions and including the word heaviness should carefully instruct their participants that this word refers to the “quality of the sensation” rather than “a weight judgement”.

## 5. Conclusion and perspectives

Our results indicate that the group III-IV muscle afferents does not contribute as a signal processed by the brain to generate the perception of effort. However, they may induce changes in the neuromuscular system, primarily by regulating the development of neuromuscular fatigue [38, 39], leading to changes in the perception of effort via their influence on the central motor command. The *effort dissociated* subgroup offers support to the corollary discharge model. Studies assigned to this group employed anaesthetics known to reduce maximal force production capacity. Consequently, participants had to increase the magnitude of their central motor command to maintain similar absolute workload, leading to an increased perception of effort. This is not seen when participants had to maintain similar relative workload (i.e., similar magnitude of the central motor command). So where does the signal(s) generating perception of effort come from? As presented in the section 4.4, various lines of evidence from exercise physiology, neuroscience and psychophysiology suggest that the perception of effort is generated by brain processing of corollary discharges. Corollary discharges are neural signals generated by premotor/motor areas of the cortex when they generate central motor commands to initiate and sustain voluntary skeletal muscle contractions [134, 135].

This meta-analysis also underscores the importance to provide clear and standardized instructions to the participants to avoid the confounding effect of other exercise-related perceptions in the ratings of perceived effort. We therefore recommend, similarly to others [14, 29, 133], that researchers and clinicians instruct and familiarize their participants to rate their perception of effort specifically by excluding other exercise-related perception(s) from their sense of effort. This is also crucial for researchers investigating the perception of effort as a psychophysiological marker of the magnitude of the motor command [e.g., 88, 90] and using muscle pain as a psychophysiological marker of feedback from group III-IV muscle afferents [e.g., 136, 137].

Investigating effort as a unique and dissociated perception is also crucial to better understand how perception of effort interacts with other exercise-related perceptions, such as pain, and influence performance and the regulation of human behavior. The dissociation between the sensory signal generating the perception of effort from other neurophysiological signals modulating this perception could help researchers and clinicians to better understand how various neurophysiological pathways [e.g., 138] or psychological factors [e.g., 1, 139] could influence this perception. It could unravel underlying mechanisms generating and regulating the perception of effort, and lead to the development of unique multidisciplinary interventions aimed at decreasing perception of effort to improve the adherence to an exercise training program [e.g., 140].

## Statement and Declarations

### Availability of data and material

All data and materials are available within the manuscript or in supplemental materials.

### Code availability

Not applicable

### Author contributions

MB, MPDLG, SM, JC and BP designed the study. MB and MPDLG performed the literature search. MB, MPDLG, CFB, JC and BP designed decision flowchart to code identified articles. MP and MPDLG performed risks of bias assessment. JS performed statistical analyses. MB, JS and BP created the figures and tables. MB and JS wrote the first draft of this manuscript. JS, SM, PR, and BP revised the first draft and final version of this manuscript. All authors approved the final manuscript.

## Acknowledgements

The authors warmly thank the librarian Marc-Olivier Croteau for his precious assistance. The authors would also like to thank the reviewers whose suggestions significantly helped us in improving the manuscript.

